# Fully automated detection of dendritic spines in 3D live cell imaging data using deep convolutional neural networks

**DOI:** 10.1101/2023.01.08.522220

**Authors:** Fabian W. Vogel, Sercan Alipek, Jens-Bastian Eppler, Jochen Triesch, Diane Bissen, Amparo Acker-Palmer, Simon Rumpel, Matthias Kaschube

**Affiliations:** Frankfurt Institute for Advanced Studies & Department of Computer Science, Goethe University Frankfurt, Ruth-Moufang-Straße 1, 60438 Frankfurt am Main, Germany; Institute for Cell Biology and Neuroscience, Goethe University Frankfurt, Max-von-Laue-Str. 15, 60438 Frankfurt am Main,Germany; Institute of Physiology, FTN, University Medical Center, Johannes Gutenberg University Mainz, Hanns-Dieter-Hüsch-Weg 19, 55128 Mainz, Germany

## Abstract

Dendritic spines are considered a morphological proxy for excitatory synapses, rendering them a target of many different lines of research. Over recent years, it has become possible to image simultaneously large numbers of dendritic spines in 3D volumes of neural tissue. In contrast, currently no automated method for spine detection exists that comes close to the detection performance reached by human experts. However, exploiting such datasets requires new tools for the fully automated detection and analysis of large numbers of spines. Here, we developed an efficient analysis pipeline to detect large numbers of dendritic spines in volumetric fluorescence imaging data. The core of our pipeline is a deep convolutional neural network, which was pretrained on a general-purpose image library, and then optimized on the spine detection task. This transfer learning approach is data efficient while achieving a high detection precision. To train and validate the model we generated a labelled dataset using five human expert annotators to account for the variability in human spine detection. The pipeline enables fully automated dendritic spine detection and reaches a near human-level detection performance. Our method for spine detection is fast, accurate and robust, and thus well suited for large-scale datasets with thousands of spines. The code is easily applicable to new datasets, achieving high detection performance, even without any retraining or adjustment of model parameters.

## Introduction

Neurons in the brain are connected via synapses to form intricate circuits, and the way information is processed and stored in these circuits depends strongly on the efficacy of synaptic connections. Most excitatory synapses in cortex are located on dendritic spines, which are small membranous protrusions from a neuron’s dendrite. A dendritic spine typically receives input from a single excitatory synapse, and the size of the dendritic spine provides a proxy for synaptic efficacy [1], [2], [3], [4], [5], [6], [7], [8]. Therefore, dendritic spines have been the focus of many lines of research, including studies on long-term potentiation (LTP) and long-term depression (LTD) of excitatory synaptic transmission [9], [10] and on neurodegenerative diseases [11], [12], [13], [14]. Advances in imaging technology enable the visualization of large populations of spines *in vivo* [15], [16], [17], [4], [18], which, potentially, could provide us with an unprecedented detailed view on the simultaneous changes of thousands of synapses, for instance, during baseline conditions [19], [20], or when an animal is involved in learning a task and forms a memory [21]. However, with the growth of such datasets, the prohibitive limitation is no longer the data acquisition, but rather the identification of spines in the recorded images. While hand-curated labeling is effective for small numbers of spines, such approach would be daunting for more recent large-scale recordings, thus requiring tools for the fully automated detection and analysis of large numbers of spines.

To extract spine positions, classical methods typically employed fixed, preselected image processing tools such as skeletonization [22], geodesic transforms [23], Scale Invariant Feature Transform (SIFT) [24] or Generalized Gradient Vector Flow (GGVF) [25]. Some studies extracted the dendritic shaft first, and then used this information to locate putative spines on it [26], [27]. However, these methods are rule-based and are very sensitive to small variations in the underlying structure of the data (e.g. different shapes of spines, imaging noise, image sizes, resolution). Their application to different datasets usually requires substantial amount of fine-tuning and expert knowledge, as such methods do not generalize well to new data obtained under different experimental conditions.

Deep convolutional neural networks (CNNs) are state-of-the-art in various fields of computer vision, including object recognition and medical image analysis. The ability of CNNs to detect complex patterns and to generalize well from specific training sets to novel data has been proven useful in a broad range of live-cell imaging applications, for instance detecting cancer [28], [29] and immune cells [30], and for cell tracking in general [31]. CNNs have also been successfully applied to data at more macroscopic scales, e.g. for analyzing the motion [32] and pose [33], [34] of individual animals and groups of animals. While some efforts have been made to use neural networks for dendritic spine detection [24], [35], [36], so far none of these methods came close to the detection performance reached by human experts.

Here, we leverage recent advances in CNNs to devise a fully automated dendritic spine detection pipeline that reaches near human-level performance (model: 0.862, human experts: 0.968) of spine detection in 3D volumes of brain imaging data of sparsely labeled neurons. We show that our method, without the need of further adjustment or training, generalizes well when applied to different datasets obtained in different laboratories using different labeling techniques and spatial resolutions. The method leverages transfer learning by adopting a CNN that was previously trained on a general visual object recognition and localization task, and retraining this network with spine imaging data. The fact that the network is pretrained drastically reduces the amount of dendritic spine data necessary for training the model. Our pipeline first recognizes and localizes putative dendritic spines in 2D images. In case these images are part of a 3D imaged volume, this information is then integrated across the entire image stack to reveal the 3D locations of dendritic spines in that volume. Our method outperforms available classical methods based on hand-crafted features as well as previous neural network based approaches. Spine detection is fast even for large numbers of spines and generalizes well to new datasets without any new training required. The code is publicly available under [37]. Thus, by providing a robust, generalizable and scalable solution, our method addresses the existing bottleneck of large-scale automated spine detection.

## Materials and methods

### Dendritic spine imaging dataset

Our main dataset for developing and validating our analysis pipeline consisted of 55 two-photon image stacks recorded in the auditory cortex of a single mouse, containing dendritic branches of GFP-expressing neurons along with their dendritic spines (Fig. 1A). This dataset was published earlier [4], [18] and details about data acquisition can be found there. The images were grayscale and had a size of 512 × 512 pixel corresponding to a linear extent of 51.2µm. The distance in z-direction between two consecutive image slices of the same stack was 0.5µm.

**Fig 1.**
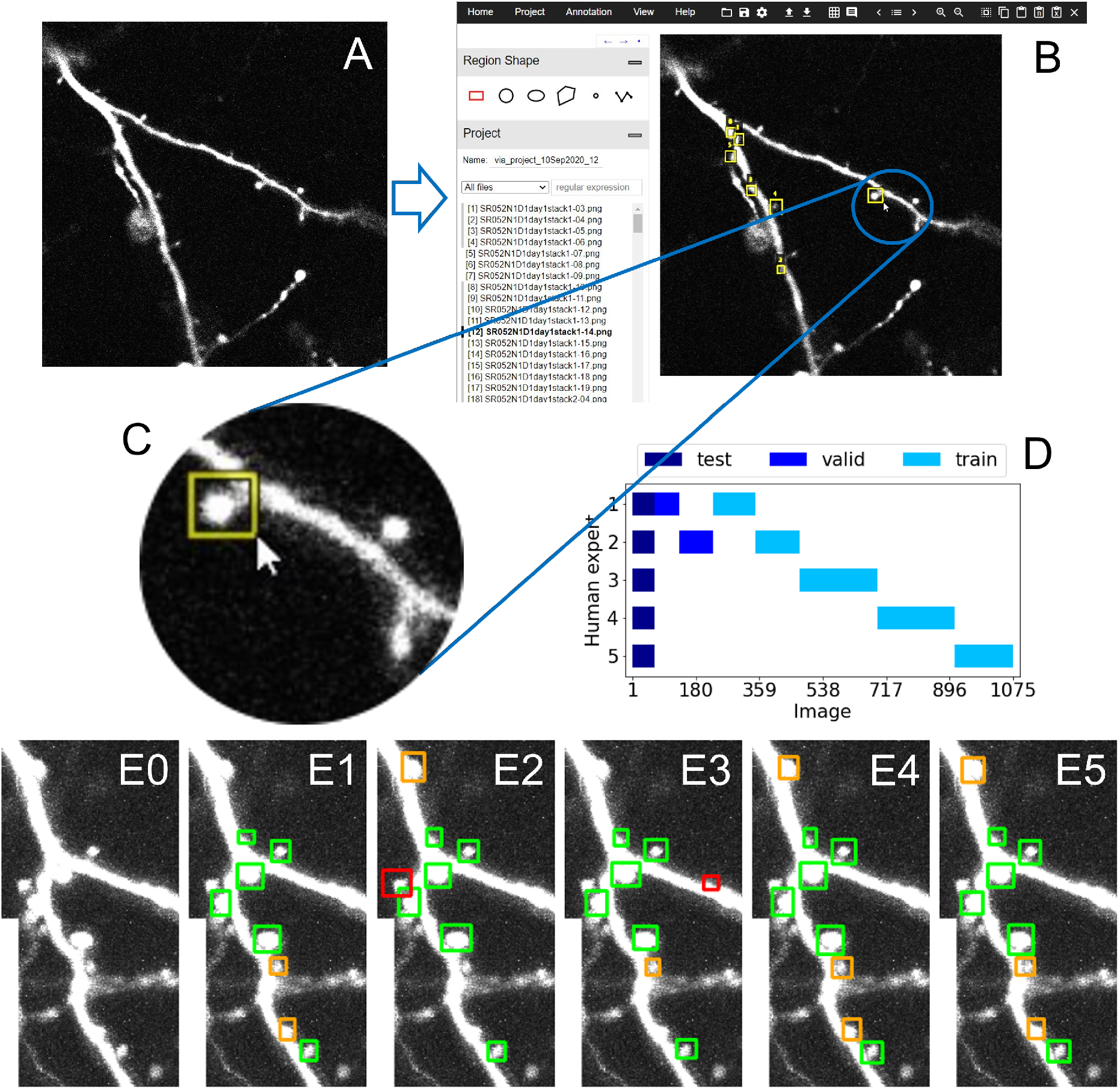
Generation of ground truth data by five human expert annotators based on prelabeling. A: Example 2D raw data image showing two dendritic branches along with several dendritic spines. B: Annotated spines from [4], [18] used as prelabels for generating the ground truth data for the present study. C: To achieve this, these prelabels are inserted into the labeling tool VIA [38], checked by our human expert annotators and adjusted further, where necessary (including deleting or creating new boxes). D: Overview showing the sets of images labeled by the five human experts and their assignment to the training, validation, and test sets. Note that the test set was labeled by all five annotators to obtain a more solid ground truth. Training sets, in contrast, were non-overlapping to maximize the overall size of the training data. E0-6: Example of an image section (E0) with the labels from all five experts (E1-5), illustrating the degree of consistency and variability across human experts. Green, labelling consistent across all experts; orange, the majority and red, the minority of experts.

Optical slices (images) that did not contain dendritic spines were excluded from the stacks, leaving 1075 slices in total that contained at least one spine each. The number of slices varied across stacks, from 5 to 68 (mean ± SD: 19.5 ± 11.5). The 55 stacks were divided into a training dataset (for training the model), a validation set (for hyperparameter optimization, see below), and a test set (for performance assessment) containing 44, 6, 5 stacks (non-overlapping) with a total of 844, 169, 62 optical slices, respectively. According to the maximal vote of human annotations (see below), these sets contained 689, 116, 55 dendritic spines, respectively. The training dataset provides the features that the network is learning in order to maximize the model performance on the validation dataset: While the network trains on the training samples, the hyperparameter choices are made with respect to an increasing validation performance. Finally, the test dataset was used to assess the model’s performance on data not included in training and validation.

### Manual labeling of data

The complete workflow of manual annotation is shown in Fig. 1. Five human expert annotators each labeled a subset of the 1075 optical slices semiautomatically using the VGG Image Annotator [38]. Each individual labeled between 228 and 282 images. A simple algorithm was used to set prelabels for putative spines in all slices, based on the labels for subsets of spines in single slices provided in [4] (Fig. 1B, C). A given dendritic spine in an image was marked by a rectangle, which the annotators were able to shift and resize to estimate the minimal bounding box containing this spine. For the majority of spines, manual labeling consisted of adjusting/correcting these prelabels. However, sometimes the insertion of new (and to a smaller degree also the elimination of existing) bounding boxes was necessary, as not all dendritic branches were labeled in the original study [4]. Note that it was necessary to include these spines and label them, as the presence of unlabeled true spines during training would confuse the network whose task is to learn to distinguish between ‘spine’ and ‘no-spine’. Also note that in several cases the structure under consideration appeared difficult to interpret, even for expert annotators (see Fig. 1 E-J for examples). To reduce such error sources, we used a two-stage process to improve the quality of labeled data: First, each annotator labeled independently. Second, their labels were reviewed by another expert using the four-eyes principle.

The data subsets that were labeled by the five annotators were chosen to be mutually non-overlapping apart from a smaller subset that was labeled by all five annotators (Fig. 1D). This overlapping region allowed us to combine the detections of the different annotators to estimate a ground truth, which we then used to evaluate the detection performances of the individual annotators as well as of our model. Specifically, we created three different ground truths from these five different human annotations: A minimal, majority, and maximal ground truth consisting of spines detected by at least one, three, or five annotators, respectively, based on an overlap score of their bounding boxes defined below. Keeping the overlapping part relatively small allowed us to distribute the individual annotations over a larger fraction of the dataset.

### Dendritic spine detection pipeline - overview

Our pipeline for automated spine detection in three-dimensional image stacks consists of several steps, shown in Fig. 2: (1) Converting stacks of images to the required format (Fig. 2A), (2) for each optical slice separately, obtaining a bounding box for every identified spine along with a confidence value of detection by applying an adequately trained Faster R-CNN model [39] (Fig. 2B), and (3) integrating this information on detected spines across all slices in the 3D stack to further improve the robustness of detection, to interpolate the spine bounding boxes across the different slices and to define the outline and location of each spine within the 3D stack (Fig. 2C). This outline, a 3D box, consists of a 3D volume confined in z by the range of optical slices a given spine was observed in (Fig. 2D) and for each optical slice confined in x and y by the bounding box. The fluorescent pixels inside this 3D volume can then be used to estimate properties of the synapse. For instance, the synaptic efficacy can be estimated by integrating the total fluorescence within this volume. In case the data consists of single image slices, step (3) is omitted.

**Fig 2.**
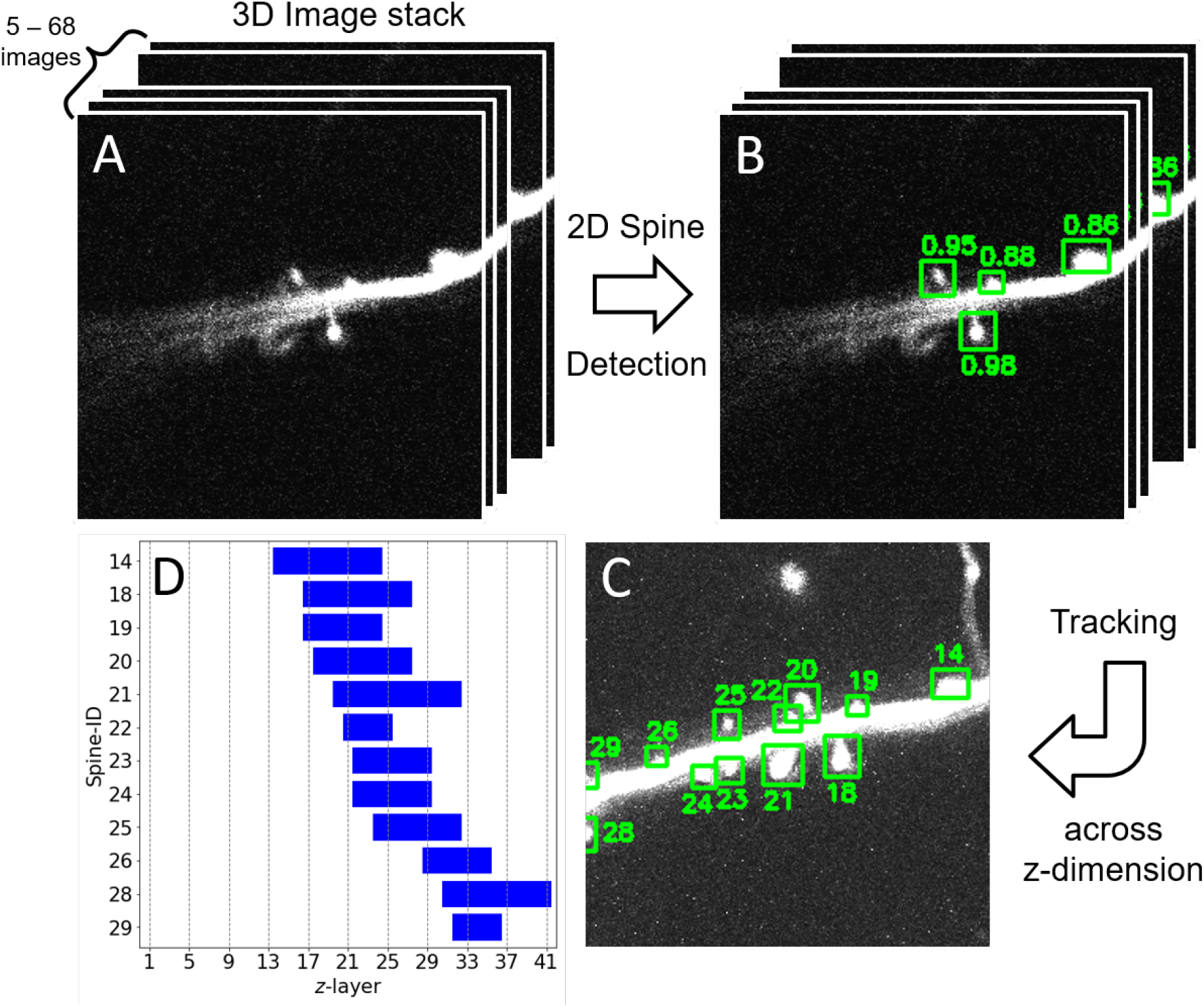
Overview of spine detection pipeline illustrated using one example image stack. A: Input to the pipeline are 3D image stacks. In *z*-direction (depth) stacks vary from 5 to 68 image slices. B: Spine detection is first carried out for each image slice, separately. For each spine a bounding box is estimated that contains the spine. Numbers indicate the estimated probability (confidence) that the corresponding box contains a spine. C: Bounding boxes are then tracked across *z*-direction. Here, the numbers denote the IDs assigned to the different spines. Bounding boxes were averaged over the *z*-direction, as boxes varied slightly between the different *z*-slices. D: The identified spines in 3D and their depth profiles. Each Spine-ID is equipped with the list of slices in *z* at which this spine was detected.

To evaluate the performance of our pipeline, we compared its detection results with the hand-labeled ground truth data, described in the *Results*.

For step (2) above, spine detection on image slices, we adopted a pretrained Faster R-CNN, with feature extractor ResNeXt-101 [40]. Its processing steps are shown in Fig. 3. The training of this network along with the choice of architecture and hyperparameter optimization is described in the next section (Section *Network training*).

**Fig 3.**
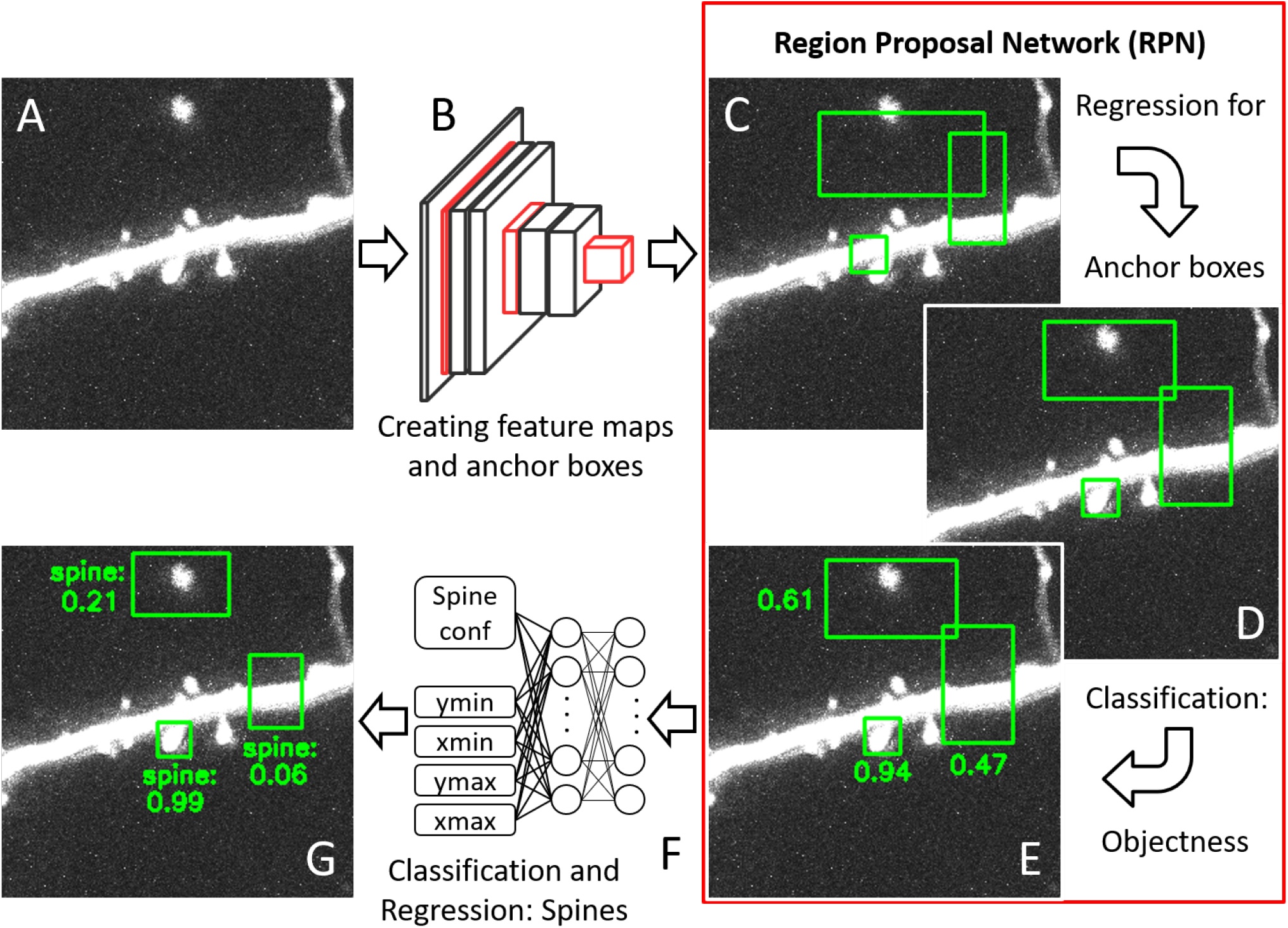
Detailed workflow of the 2D spine detection part (Fig. 2B) using a region proposal network (RPN). A: A part of a raw image slice with several spines along a dendritic branch. B: A sequence of multiple convolutional layers transform this image into sets of feature maps capturing relevant geometric properties in the image. C: Anchor boxes of varying size and aspect ratio are generated (stride of eight, twelve anchor boxes per location). For the sake of clarity only three examples are shown. D: The RPN then refines the location of these anchor boxes using regression. E: For each box, the RPN calculates the probability (indicated by the numbers) that it contains an object (classification for objectness). F: After region of interest (ROI) pooling, fully connected layers feed into i) one output neuron representing the confidence of a spine being inside a given anchor box (classification for presence of spine) and ii) four output neurons estimating the top left and bottom right corner of this anchor box (via a regression step), G: Final output image with bounding boxes around the detected spines along with confidence values (max, 1; min, 0).

### Network training

Testing a variety of neural network architectures combined with different feature extractors, we found that using the Faster R-CNN [39] with feature extractor ResNeXt-101 [40] as our base network provided promising results and, thus, we focus on this architecture in the following. The model was pretrained on the MS COCO dataset [41] for visual object recognition and afterwards trained on our spine dataset. Pretraining established the model’s general ability to detect a broad range of visual objects, along with the necessary sensitivities for visual features on various spatial scales and levels of complexity. Adapting such a pretrained model enabled us to cope with a relatively small amount of training data (and training time), while achieving a high detection performance on tests with novel spine imaging data.

The network consists of two main parts: First, the Region Proposal Network (RPN), which generates an array of candidate bounding boxes (N=50,700), the so called anchors (Fig. 3C). Anchor positions are further refined by regression and for each anchor a confidence score is computed expressing whether it contains an object or not (Fig. 3D, E). Applying non-maximum suppression (using the intersection over union (IoU) score with threshold 0.7) and based on the highest confidence scores, a subset of at most *N* ^RPN^ = 1000 of these boxes, the so called region proposals, are selected and fed into the second part of the network, the Box Classifier (BoxC) (Fig. 3F). This part performs Region of Interest (RoI) Pooling, refinement, and classification through a set of fully connected layers to obtain a reduced and final list of *N* ^BoxC^ = 80 bounding boxes with their confidence of containing a dendritic spine (Fig. 3G). Both networks were retrained jointly.

Training the first part, the RPN, involves a comparison between all the region proposals (i.e. the potential anchor boxes) and the ground truth boxes given by the labeled data. For each ground truth box the region proposal with the highest *IoU* overlap gets assigned the label 1, if this overlap exceeds a threshold *IoU* ≥ 0.7. The remaining boxes with an *IoU* overlap below 0.3 for all ground truth boxes are not sufficiently close to any of the ground truth boxes, and they receive the label 0. All other boxes, which are assigned to neither of these labels, do not contribute to the training objective; thus, are not included in the loss calculation. This procedure is in line with the general training scheme used for the Faster R-CNN [39]. The loss function then comprises two contributions: First, the objectness loss ℒ_*obj*_ penalizing if the labels (object vs. non-object) of the anchors are correct, and second, the regression loss ℒ_*regr*_ that penalizes deviations in the spatial position and size of anchor boxes. Both losses are normalized by the corresponding number of proposal regions. Their weighted sum yields the overall loss ℒ^*RPN*^ (ℒ_*regr*_ is weighted by a factor *α* = 2 relative to ℒ _*obj*_ to prioritize regression over classification).Training the second part involves training the box classifier, whose structure is similar to the structure of the RPN, but placed at a different location within the network. Fig. 3C-E shows the location of the RPN and Fig. 3F the location of the box classifier. The loss of the Box Classifier is denoted with ℒ^BoxC^ and is calculated similar to the loss ℒ^RPN^ of the RPN. However, the mapping between predictions and ground truths differs slightly as the set of predictions of the Box Classifier consists of only *N* ^BoxC^ = 80 refined boxes, as opposed to the *N* ^RPN^ = 1000 anchor boxes considered for the RPN loss, above, requiring an adjustment of the normalization constants. All other parts of the ℒ ^BoxC^ calculation are analogous to that of ℒ^RPN^.

Finally, to avoid large weights inside the neural network, an additional L2 regularization loss ℒ^regu^ is added with a factor of *λ* = 3 · 10^*−*6^. The combined loss function then reads:

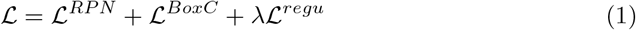

We trained the model using a batch size of 1 and gradient descent with momentum of 0.6 and learning rate of *γ* = 1 · 10^*−*3^. By visual inspection of the approximate spine sizes in the labeled images we chose the following parameters for the anchor generator:Scales of 8, strides of 4, 8, 16, 32, 64 pixel, and aspect ratios of 0.5, 1.0, 2.0. We used in-place data augmentation to increase the complexity of the dataset without increasing its size. To this end, we randomly applied a right-turning 90 degree rotation, vertical and horizontal flips, leading to eight different orientations of the original image, followed by adding Gaussian white noise and Gaussian blur, each with probability 0.5. In total, the network was trained for 14348 iterations (17 epochs). To find optimal values for the hyperparameters momentum, learning rate, weight decay and the in-place data augmentation, we conducted a two-staged grid search, starting with a coarse grid followed by a finer sampling within the hyperparameter space.

All computations related to CNNs were conducted in Pytorch [42]. Training for 17 epochs took slightly less than one hour on an Nvidia RTX A6000 GPU. Our model took on average 0.053 seconds to make predictions on one optical slice. As it will always return 80 bounding boxes for spines, no matter how low the confidence score is, the detection time does not depend on the number of spines visible. Tracking one detection into the next frame took 0.01 seconds, which is negligible in comparison to the detection time. In all images of our dataset, the number of spines never exceeded 26. In the case that more than 80 spines are visible, the image can be split up into *n* parts so that each part only contains less than 80 spines. This will multiply the detection time by a factor of *n*. Alternatively this upper bound can be manually increased (requiring retraining of the model).

### Evaluation metric 2D

The evaluation of the accuracy of spine detection is based on the overlap between the detected and the ground truth boxes. To compute this overlap we defined the **Intersection over Minimum *IoM***_*xy*_ for image slice *i* in the *xy*-plane as

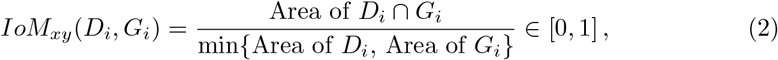

where *D*_*i*_ is a detection box and *G*_*i*_ a ground truth box in this image slice. We found that this metric achieves more accurate results compared to the more commonly used metric Intersection over Union (*IoU*). This appears to be due to the fact that dendritic spines are small compared to the entire image and so slight positional differences in the estimated bounding boxes can significantly affect the value of the *IoU*. Note that this issue is not solved simply by reducing the detection threshold of the *IoU*, as this produces more false positives. Fig. 4A-C and the schematic Eq. (6) illustrate the difference between the *IoU* and the *IoM*_*xy*_. Fig. 4D-H illustrates the *IoU* and *IoM*_*xy*_ scores using hand-crafted bounding boxes that were chosen to emphasize the differences between these measures. In our data we found that using *IoM*_*xy*_ typically yielded more robust results. In few cases the *IoM*_*xy*_ can be misleading if the bounding box is considerably larger than the ground truth. Thus, we introduced an area threshold to remove all bounding boxes exceeding a specific size of 2000 pixel (20*µm*^2^). This value is chosen according to the observed statistical spine properties: Most spines had an area between 100 and 700 pixel (1− 7µm^2^), the largest spine in our labeled dataset achieved an area of 1300 pixel (13µm^2^). As our model is trained to minimize the loss which depends on the exact position of the bounding box, such cases are extremely rare. We thus conclude that the metric ***IoM***_*xy*_ is better suited for the purpose of spine detection as it represents the quality of human annotations and detections more accurately.

**Fig 4.**
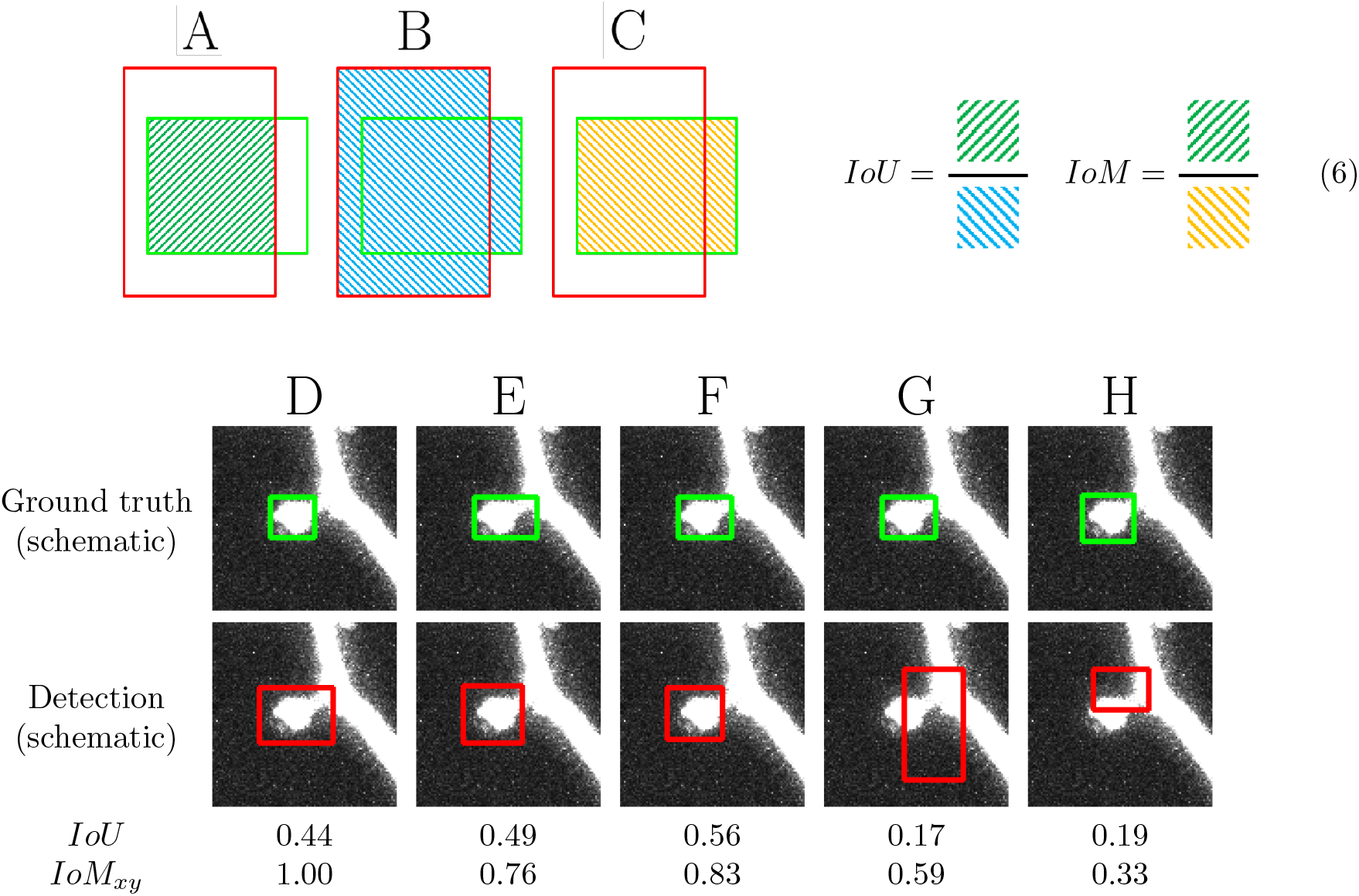
Intersection over Minimum (*IoM*) is better suited than Intersection over Union (*IoU*) for evaluating spine detection accuracy. A-C: Given a ground truth box (green) and a detection box (red) their overlap can be defined by the *IoU* or the *IoM*, both of which are defined in the schematic equation (right, Eq. (6)), where the squared symbols colored in green, blue, and yellow denote the areas of intersection (A), union (B) and minimum area (C), respectively. Note that the two equations differ only in their denominator. D-H: Illustration of the difference between *IoU* and *IoM*_*xy*_. For fairly accurate detections (D-F), the *IoU* can be quite low, while the *IoM*_*xy*_ appears to reflect better the detection accuracy. For seemingly false detections (G, H) the *IoM*_*xy*_ is typically low (H), but can be inaccurate in rare cases (G). Note that the bounding boxes depicted here are drawn manually to illustrate the difference between *IoM* and *IoU* .

### Integration across imaging depth

When applied to 2D data (single images), our method estimates dendritic spines as described up to this point. When a 3D image stack is provided, our tool integrates the information across depth, as described in the following.

If an image stack is provided, we use a tracking algorithm to combine detections across layers within the stack. This improves the robustness of detection (in the case a misdetection occurred in a few layers), and allows us to extract the *x*-, *y*- and *z*-coordinates of identified spines.

The algorithm proceeds as follows. First, the model yields a list of bounding boxes, each equipped with a confidence score (Fig. 3G). Next, the number of model predictions is reduced to only keep the relevant detections: Only boxes that have a confidence greater than the commonly used prediction probability threshold of 0.5 are considered as real detections. If multiple detections with a pairwise *IoM*_*xy*_ of more than 0.5 exist, all those detections except the one with the highest detection probability will be removed.

In the last step, we devised a simple tracking algorithm that converts this set of remaining detections from the different optical slices in a stack into a list of identified spines for that stack. It starts with iterating over the *z*-axis from top to bottom. In each iteration, the identified spines from the current optical slice are compared to those in the next slice, using the *IoM*_*xy*_-score. Pairs of identified spines with the highest *IoM*_*xy*_-scores are considered to belong to the same spine, if this score exceeds the threshold of 0.5. After connecting all spines, some spines detected in the next slice may be left without any assignment, in which case this spine was registered as a new spine. To account for possible missed detections in individual slices, the tracking of a given spine only stopped after two consecutive missing detections.

### Evaluation metric 3D

Once spines have been tracked across depth in an image stack, we can define a bounding box in 3D along with a measure to compare such 3D boxes between model prediction and ground truth and between annotators and ground truth.

Each 3D detection box 𝒟 consists of multiple boxes 𝒟_*i*_ which correspond to the box in image slice *i*. The values of *i* are taken from the interval 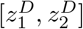representing theboundaries of the detection 𝒟 in the *z*-dimension. Analogously, each ground truth box 𝒢 is assembled by 2D boxes 𝒢_*i*_ for 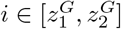. The intervals are returned by the tracking algorithm above (see also Fig. 2.D). Analogously to the *IoM*_*xy*_, we defined the *IoM*_*z*_ representing the percentage overlap in *z*-direction of these intervals:

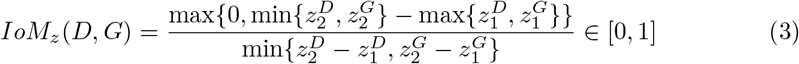

Further, we computed the *IoM*_*xy*_ for 3D detection and ground truth boxes 𝒟, 𝒢 by averaging in *z*-direction over all associated 2D boxes:

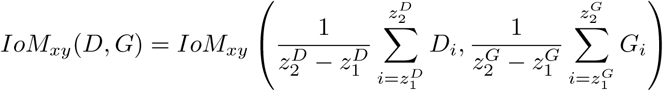

These two scores, *IoM*_*xy*_ and *IoM*_*z*_, were then combined using the *F*_0.5_-score. The parameter *β* = 0.5 is chosen to increase the importance of *IoM*_*xy*_ relative to *IoM*_*z*_. The final *IoM* -score given a 3D detection 𝒟 and a 3D ground truth label 𝒢 is then defined as:

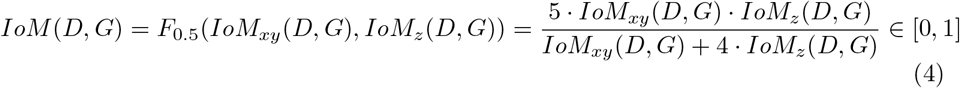

We defined true positives (TP), false positives (FP) and false negatives (FN) similar to the common definition but using the *IoM* instead of the *IoU* to decide if a detection is correct or not. The specific definitions are as follows:

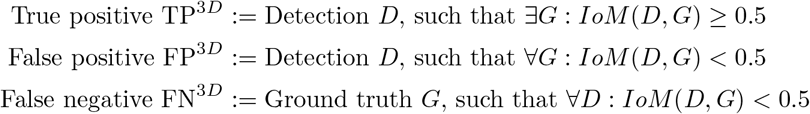

Finally, to evaluate the overall detection performance, we defined the 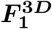**-score**, which is similar to the standard *F*_1_-score, but with the above definitions of TP^3*D*^, FP^3*D*^ and FN^3*D*^. Given a list of detections 𝒟 = 𝒟^(*j*)^} and a list of ground truth labels 𝒢 = {𝒢^(*k*)^}, this score is given by

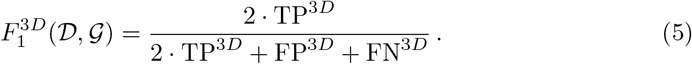

The 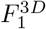-score was used both for comparing our model in 3D against ground truth and for comparing the detections of individual annotators against ground truth.

## Results

Spine labels varied slightly across the five expert annotators (Fig. 1 E1-E5), suggesting that for some cases it was difficult to decide between spine or no spine, even for a trained human expert. To account for this ambiguity we considered three alternative definitions of ‘ground truth’ data, against which we evaluated the performance of both our individual experts and our model. The ‘minimal ground truth’ comprised all spines that were identified by at least one of the experts, while the ‘majority ground truth’ comprised all spines identified by at least three experts, and ‘maximal ground truth’ the set of spines all five experts agreed on. Our assessment of performance focused mostly on the majority ground truth, as we expect it to reflect the most accurate estimate of the true set of spines.

Before investigating the overall performance of our model on the three-dimensional data stacks, we first examined the accuracy of spine detection when applying the model to individual two-dimensional image slices. After hyperparameter optimization the model’s performance on the validation dataset, based on the standard *F*_1_-score and averaged over four runs, reached 0.802 ± 0.012 (mean ± SEM). In comparison, the performance on the test dataset was slightly lower for all three types of ground truths: 0.747 ± 0.032 (minimal), 0.758 ± 0.028 (majority) and 0.741 ± 0.025 (maximal). A gap between validation and test performance is expected given the relative small number of image stacks within these two datasets and may in parts also reflect subtle differences across stacks regarding biological properties or the experimental conditions.

To leverage the full potential of the volumetric spine imaging datasets, we added a tracking scheme across the layers of the stack to integrate the detection information across the different image slices within the stack. This strategy allowed us to compensate for some of the detection failures or inaccuracies in the individual slices, resulting in overall more robust spine detection performances. To quantify this performance, we defined a novel three-dimensional performance metric, the 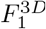-score (see Methods), and applied it to detections made using the testdata (which was neither used for training nor hyperparameter optimization). Fig. 5 summarizes the results for the detection performance of our model in three dimensions for the three types of ground truth, and in comparison to the performance achieved by the five human expert annotators. For the model, the 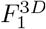-scores averaged over four randomly initialized training runs were 0.836 ± 0.025 (minimal), 0.843 ± 0.0082 (maximal) and0.862 ± 0.0087 (majority). Note that these performances were achieved using the hyperparameters, for which we obtained the maximal *F*_1_-score for the two-dimensional detection in the image slices of the validation dataset. We noticed, however, that when training the model longer, this typically resulted in even higher 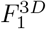-scores on the test dataset. For instance, for the majority ground truth, our model achieved a 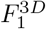-score of 0.903 when trained over 26 instead of 17 epochs (orange dashed line in Fig. 5).

**Fig 5.**
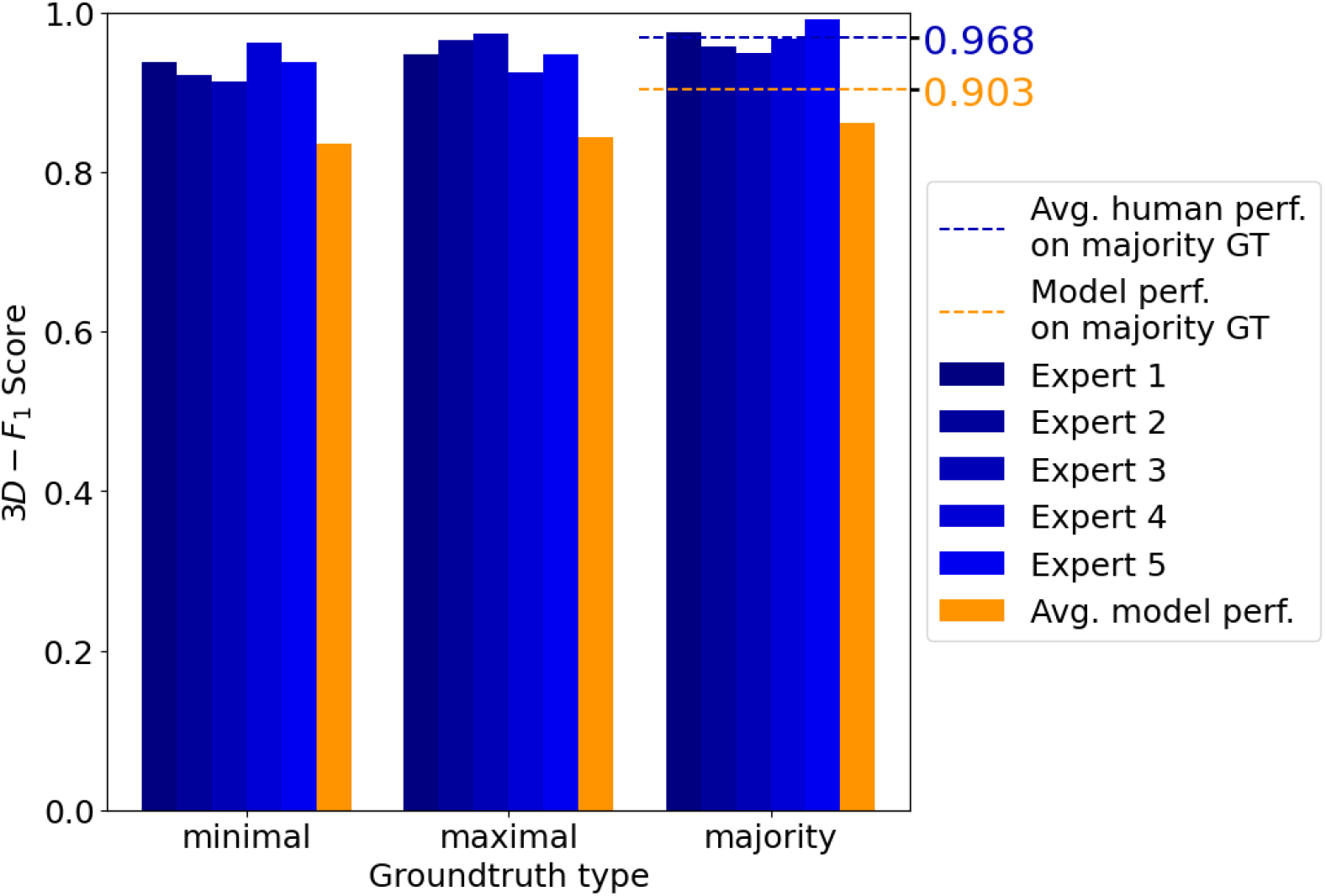
Automated spine detection reaches near human-level performance. Detection performance 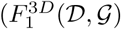, see section *Evaluation metric 3D*) of our model compared to the performance achieved by five human experts. To assess performance, detection is compared with three different types of ground truth (GT): minimal (left),maximal (middle) and majority (right), defined by the set of spines detected by at least one, all five and at least three experts, respectively. The model performance is averaged over four randomly initialized training runs. Using the hyperparameters with the best two-dimensional *F*_1_-score on the validation set yields a model performance of 0.836 ± 0.025 (minimal), 0.843 ± 0.0082 (maximal) and 0.862 ± 0.0087 (majority ground truth) (orange bars). Even slightly higher test results were achieved for longer training times (26 vs. 17 epochs; orange dashed line, 0.903).

In comparison to the model, the human annotators reached slightly higher 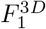-scores with an average value of 0.968 (blue dashed line). Fig. 6 depicts examples of correctly detected spines along with a few examples of misdetections by the model.Examples of the latter include, for instance, cases of close proximity, which the model counts as a single spine due to the high overlap of two bounding boxes. Thus, overall, our model achieved a detection performance that came close to that of trained human experts. However, once trained, the time it took the model to process the data was substantially shorter compared with human experts. While our well-trained human annotators reached rates of 12 seconds per frame (for frames containing only few spines), the model processed a frame in 0.053 seconds (independent of spine number) using an Nvidia RTX A6000. This fast automated detection of spines makes the model readily applicable to large datasets.

**Fig 6.**
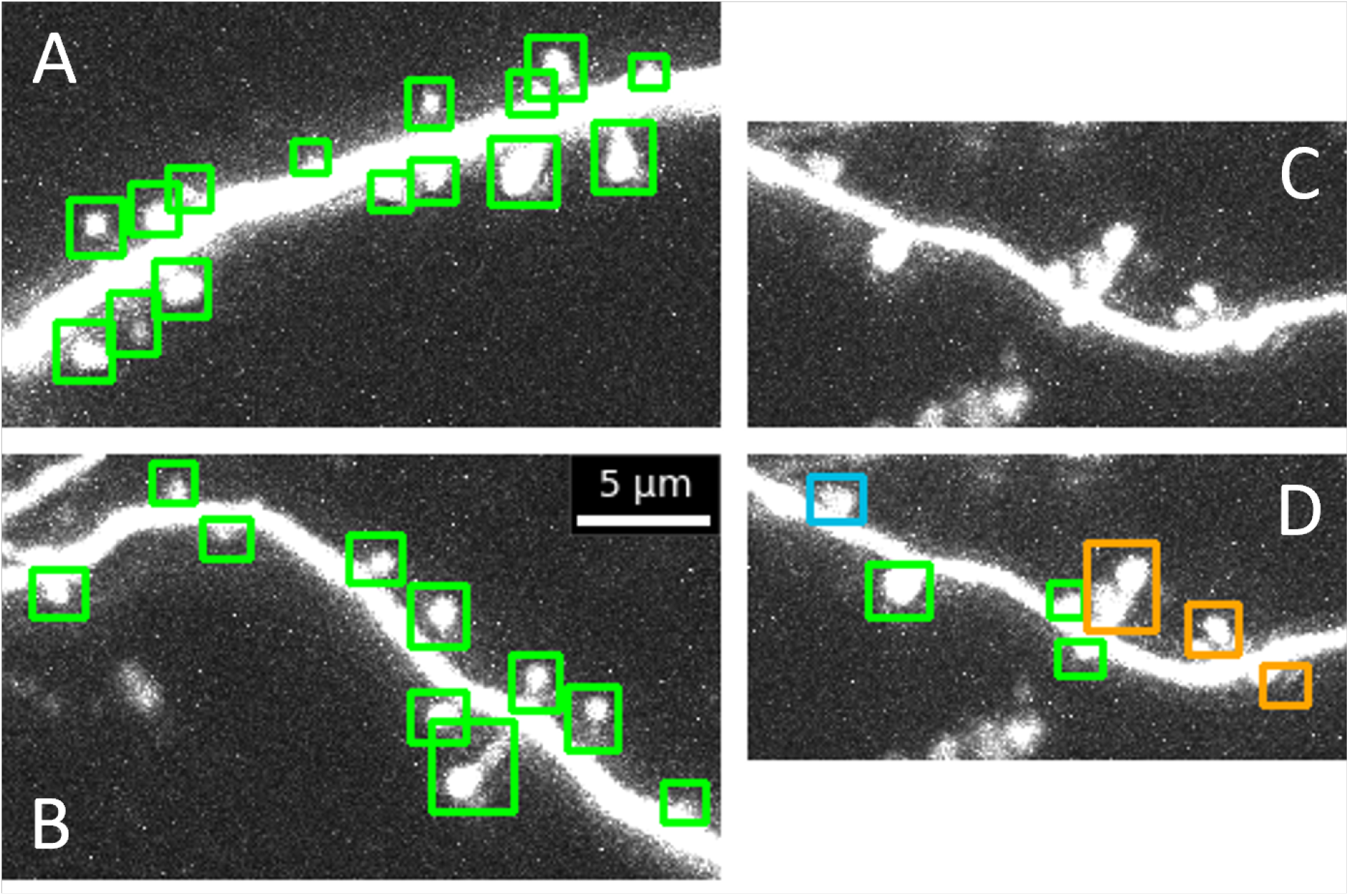
Examples of correctly identified spines and detection errors in novel data (from a mouse that was not used for training/validation/testing of the model). Images and bounding boxes are shown as maximum intensity projections (MIP) (which for the bounding boxes is equivalent to taking a 2D slice through the 3D box identified). A, B: Examples of correctly detected spines, C: An example of an image section in which spine detection is particularly challenging. D: Model results: some spines are correctly detected (green), but one is missed (blue) and several nearby spines are merged into a single detection (orange). Note that examples like these are rare in our data, but shown here to illustrate potential limitations of our model.

Next, to test the model’s robustness and generalizability, we applied it (without any further adjustment) to novel datasets. First, the model was applied to image stacks obtained from a different mouse (same experimental setup) that was neither used for training nor validation of the model. The example shown in Fig. 7A illustrates the overall high accuracy of the model, i.e. its detection of spines and their delineation with bounding boxes, and highlights the robustness of the model against animal-to-animal variability, noise and various background structures. Notably, the model was neither confused by axons (marked in red in Fig. 7A for display purposes) nor by other high fluorescence structures of similar size (e.g. the many puncta and protrusion-like structures in Fig. 7A). Validating the model detections by a human expert yielded a *F*_1_-score of 0.969 with a precision of 0.969. These results suggest that the detection method generalizes well to novel datasets from different specimen obtained under similar experimental conditions.

**Fig 7.**
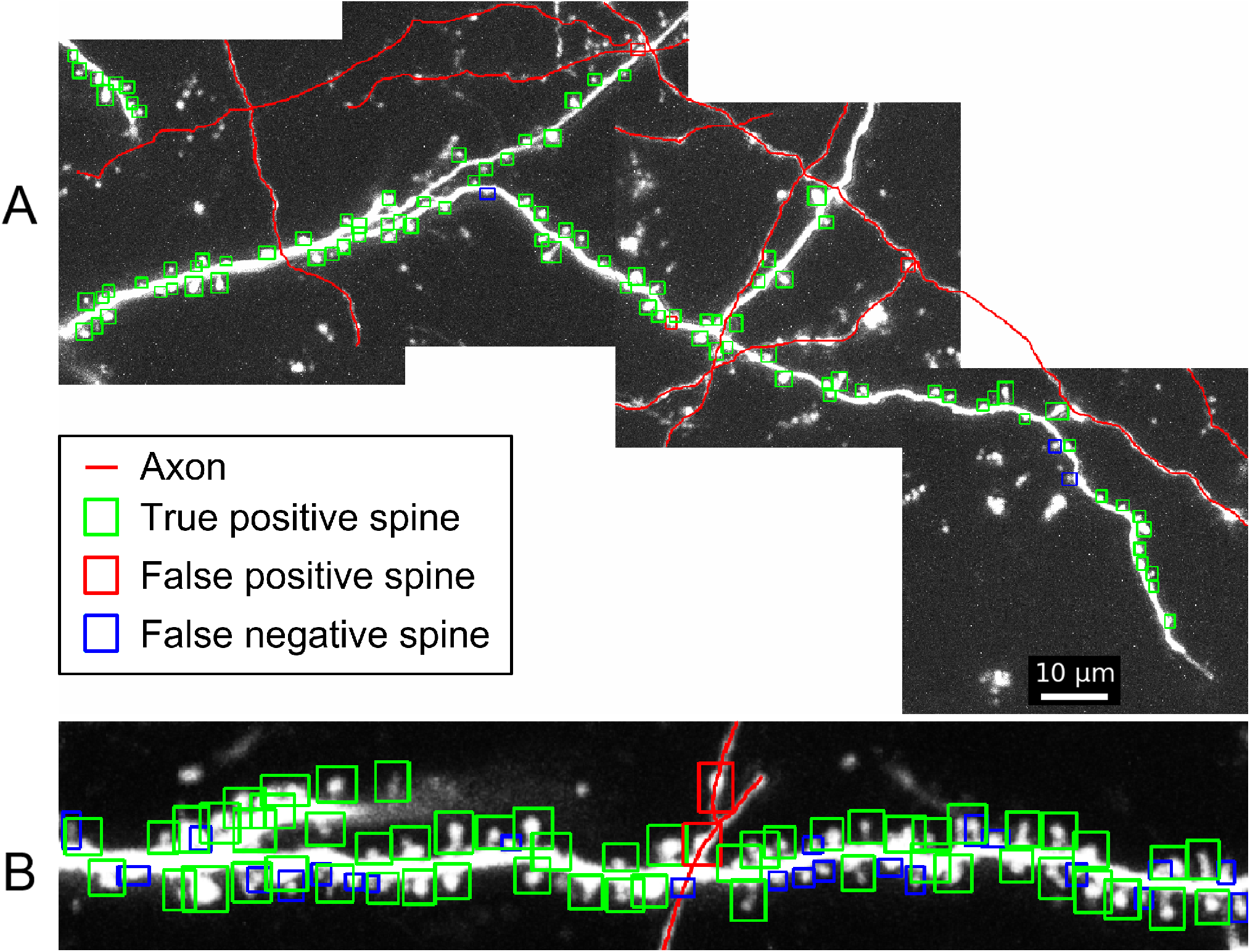
Large-scale detection of spines and its generalization to different datasets. A: Application of detection pipeline to new data (same experiments as in the previous figures, but for a different animal not used for training, validation or testing of the model). Green boxes indicate correctly identified spines, red boxes falsely identified spines and blue boxes missing spines. Red lines mark axons (highlighted manually). Note that most spines are correctly detected, despite significant background noise. B: Spines detected in a different system imaged in a different laboratory (organotypic slice cultures from hippocampus).

Second, as a more challenging test, in Fig. 7B we applied the model (without any further adjustment) to an entirely different spine imaging dataset that was obtained by a different laboratory in a different system (organotypic slice cultures from hippocampus vs. auditory cortex *in vivo*) using a different GFP label and a different microscope system with a different z-resolution (0.66*µm* vs. 0.5*µm*). As can be seen in Fig. 7B almost all spines were identified correctly by our model, achieving a *F*_1_-score of 0.815 with a precision of 0.965 on this completely new dataset. Most of the missed spines were close to another detected spine in the *xy*-plane, and could potentially be distinguished when processing different *z*-layers. This suggests that our model can achieve a near human-level performance in spine detection even in entirely novel datasets.

## Discussion

We developed an efficient analysis pipeline to detect large numbers of dendritic spines – a proxy for excitatory synaptic connections – in fluorescence imaging data. The core of our pipeline is a deep convolutional neural network, which is pretrained on the task of object detection, and which we adjusted and trained specifically to detect spines in 2D images of sparsely fluorescent-labeled neurons. In case the data consists of 3D image stacks the detection algorithm is combined with an algorithm for tracking the identified spines across depth, which further improves the overall robustness of spine identification and enables us to reconstruct the outlines and positions of spines in 3D volumes of brain tissue. To train the model we used labeled data curated by five independent human expert annotators and we combined the overlapping parts of these five sets of annotations to estimate ground truth labeled data, which we used to test the detection performance of both our model and the individual annotators. For evaluation in 3D we devised the 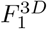-score, an extension of the widely used *F*_1_-score, well suited for comparing the 3D bounding boxes of detected spines with those in the ground truth data. We found that our method is fast and accurate, achieving near human-level performance on the ground truth test data. Most importantly, our method achieved accurate results also when applied to novel datasets that were obtained by a different laboratory in a different neural tissue, and without the need for any adjustment or re-training of the model, suggesting that it could be applicable with similar performance to a broad range of spine imaging data. The code for our method is available under [37].

When generating labeled data for training our network, we deliberately chose the labeled sets to be largely non-overlapping among our five annotators, while overlap was enforced in a subset of the data, allowing us to accurately estimate ground truth for testing and performance evaluation using that subset. This strategy was chosen in order to efficiently use the available resources for labelling. For network training it was advantageous to keep the other labeled parts non-overlapping, as this allowed the network during training to sample from a much broader set of different spines than would have been possible if all labeled sets were overlapping among the five annotators (using the same total number of annotations). Similarly, also the validation dataset, on which the model hyperparameters were optimized, consisted of non-overlapping labeled data. While it is expected that the labeling in these non-overlapping parts of the datasets are more prone to noise and personal biases in the annotation procedure, this seems more than compensated by the larger number and diversity of labeled spines.

Moreover, incorporating such biases during training may very well improve the overall robustness and generalizability of our spine detection method, which is particularly useful when applying it to novel types of datasets.

Several previous studies have used neural networks for automated spine detection [24], [36] and [35]. The method by [36] first extracts features and then trains a neural network on these features to detect spines, instead of applying a neural network directly to raw images. While this method worked fairly well for the particular dataset used in this study, due to the predefined features the model appears more rigid. Moreover, the results were not compared with labels from multiple annotators. For these limitations, it is not clear whether this model generalizes well to other datasets. The approach by [24] has the advantage of traceability of the detection process. However, it was already outperformed by the other methods. The study [35] uses a method more similar to ours, but reduces the object detection task to detecting the center of spines, which neglects its shape. Nevertheless, our approach outperforms this model in terms of detection performance and speed, and we show generalizability to new, distinct datasets. While these previous methods apply neural networks to the task of spine detection, none of them does so by exploiting information across three dimensions. The study [35] uses 3D images but then calculates the maximum intensity projection. With our tool it is possible to flexibly switch between fast detections using maximum intensity projection and detailed three-dimensional tracking of dendritic spines. The recent work by [43] introduces a comprehensive dendritic spine analysis GUI, which contains a CNN-based spine detection pipeline. The detection model is trained from scratch using large amounts of labeled training data, achieving a satisfying performance on the data sets used for model training. Our model, owing to the fact that it is based on a pre-trained general purpose detection network, not only requires fewer labeled data for model training, but also generalizes well when applied to novel data not used for training, even across different brain areas, imaging conditions and laboratories.

In summary, we devised a new method for spine detection that is fast, accurate and robust, and thus well suited for large datasets with thousands of spines. The method achieves a high detection performance, even when being applied to novel datasets obtained under different experimental conditions. Moreover, the method is flexible and can be further improved by anyone using additional suitable data for training.

## Acknowledgments

We thank Dominik Aschauer, Noelia Mateu Fernández and Takahiro Noda for annotation of spine data. This work was supported by the research grants DFG SPP 2041 “Computational Connectomics” (J.T., S.R., M.K.), DFG CRC 1080 “Neural Homeostasis” (A.A.P., D.B., S.R.), DFG/ANR project no. 431393205 (to S.R.), and DFG DIP “Neurobiology of Forgetting” (S.R. and M.K.).

## Notes

https://github.com/SaILaIDiN/Spine-Detection-with-CNNs

### Competing Interest Statement

The authors have declared no competing interest.

